# MiRNA-21-5p and MiRNA-125-5p Serve as Biomarkers for Diagnosis and Prognosis of Gastric Cancer

**DOI:** 10.1101/2024.08.02.606443

**Authors:** Xin Zhang, Yao Xiang, Junpu Wang

## Abstract

**Background:** Gastric cancer is one of the most common diseases worldwide. MicroRNAs (miRNAs) are small non-coding RNAs, typically 19-25 nucleotides in length, which represent potential targets for therapeutic intervention.

**Methods and Results:** In this study, we performed a data mining analysis for gastric cancer (GC) by integrating differentially expressed genes obtained from the Gene Expression Omnibus (GEO) database with data from The Cancer Genome Atlas (TCGA) database. We compared the expression levels of miRNA-125-5p and miRNA-21-5p in cancer cells and GES-1 cells. Our analysis revealed up-regulation of these two miRNAs in both GEO and TCGA datasets, which was further validated by quantitative polymerase chain reaction (qPCR). Additionally, survival analysis, risk score analysis, and downstream analyses, including functional enrichment analysis, target gene prediction, and protein-protein interaction analysis, were conducted.

**Conclusion:** The expression levels of miRNA-125-5p and miRNA-21-5p are upregulated in gastric cancer cells, which contribute to the diagnosis of gastric cancer and serve as novel biomarkers.

## Background

Due to early treatment of Helicobacter pylori [1-3] and advancements in medical care[4, 5], the age-standardized death rate of gastric cancer (GC) has significantly decreased over the past three decades in China[6]. However, despite these improvements, GC still presents a grim prognosis and remains the third leading cause of cancer-related deaths, following lung and liver cancer[3], partly due to late-stage diagnosis and the lack of available targeted therapy. The vast majority of gastric cancers are adenocarcinomas, originating from the mucosa or glands of the stomach’s superficial layer. Therefore, we have a keen interest in this type of GC and are eager to identify essential biomarkers and precise targets for its development and progression.

MicroRNAs (miRNAs) are short single-stranded noncoding endogenous RNAs, typically 19–25 nt in length, that regulate post-transcriptional silencing of target genes[7]. A single miRNA can target hundreds of mRNAs and it can also influence the expression of many genes [8]. Moreover, there is a wealth of evidence indicating that miRNAs can be aberrantly expressed in cancer, suggesting that they can be either oncogenic or cancer-suppressive [9]. Additionally, miRNA expression levels vary among different tissues and organs and are associated with overall patient survival, tumor stage, and the development of metastases and recurrences[10]. In clinical treatment, miRNAs hold promise as biomarkers for accurately predicting patient outcomes [11]. Thus, we aim to analyze the data using bioinformatics methods.

Bioinformatics, which combines molecular biology and information technology, has become one of the most active fields in life science. It encompasses the use of statistical, mathematical, and computational methods for processing and analyzing data[12]. In our present study, we conducted data mining analysis for GC by integrating the differentially expressed genes acquired from the Gene Expression Omnibus (GEO) database into The Cancer Genome Atlas (TCGA) database, and subsequently identified new biomarkers through bioinformatics analysis.

The Gene Expression Omnibus (GEO) Database allows users to download and analyze the data stored within it, including high-throughput functional genomics data generated from microarray-based and sequence-based technologies[13, 14]. The Cancer Genome Atlas (TCGA) database, supported by the National Institutes of Health, contains a wealth of renowned public databases containing genetic information[14].

In our study, we used bioinformatics methods and techniques to analyze and integrate miRNA expression data of GC from GEO and TCGA. Subsequently, we identified two specific miRNAs (miRNA-21-5p, miRNA-125b-5p) in GC. Based on these miRNAs, we further investigated functional enrichment analysis, protein-protein interaction, competing endogenous RNA (ceRNA) network, and survival analysis. To verify our results, we conducted PCR detection in cancer cells and GES-1 cells. This study may provide deeper insights into the transcriptomic and functional features of GC, and suggest potential therapeutic targets and biomarkers for GC.

## Methods

### 1. Microarray studies, datasets and clinical sample characteristics from GEO data repository

In our study, we compared the miRNA expression profiles from GEO and TCGA datasets. Studies were considered eligible for further analysis based on the following criteria: (1) Studies providing information about the technology and platform utilized. (2) Studies including normal groups as controls. (3) Studies including tissue samples from patients with adenocarcinomas. Following these criteria, we downloaded 433 GC tissues and 41 adjacent normal tissues from TCGA, along with two datasets from GEO. One dataset, GSE93415, comprised 20 primary GC tissues and adjacent normal tissues, while the other, GSE106817, included blood samples from 115 GC patients and 100 healthy individuals. Principal component analysis (PCA) was conducted for dimensionality reduction and quality control. Sample descriptions are provided in Table 1, and clinical information and sample sizes for the TCGA dataset are detailed in Table 2. Our workflow for the bioinformatics analysis of publicly available datasets from both GEO and TCGA databases is illustrated in Figure 1.

**Table 1.**
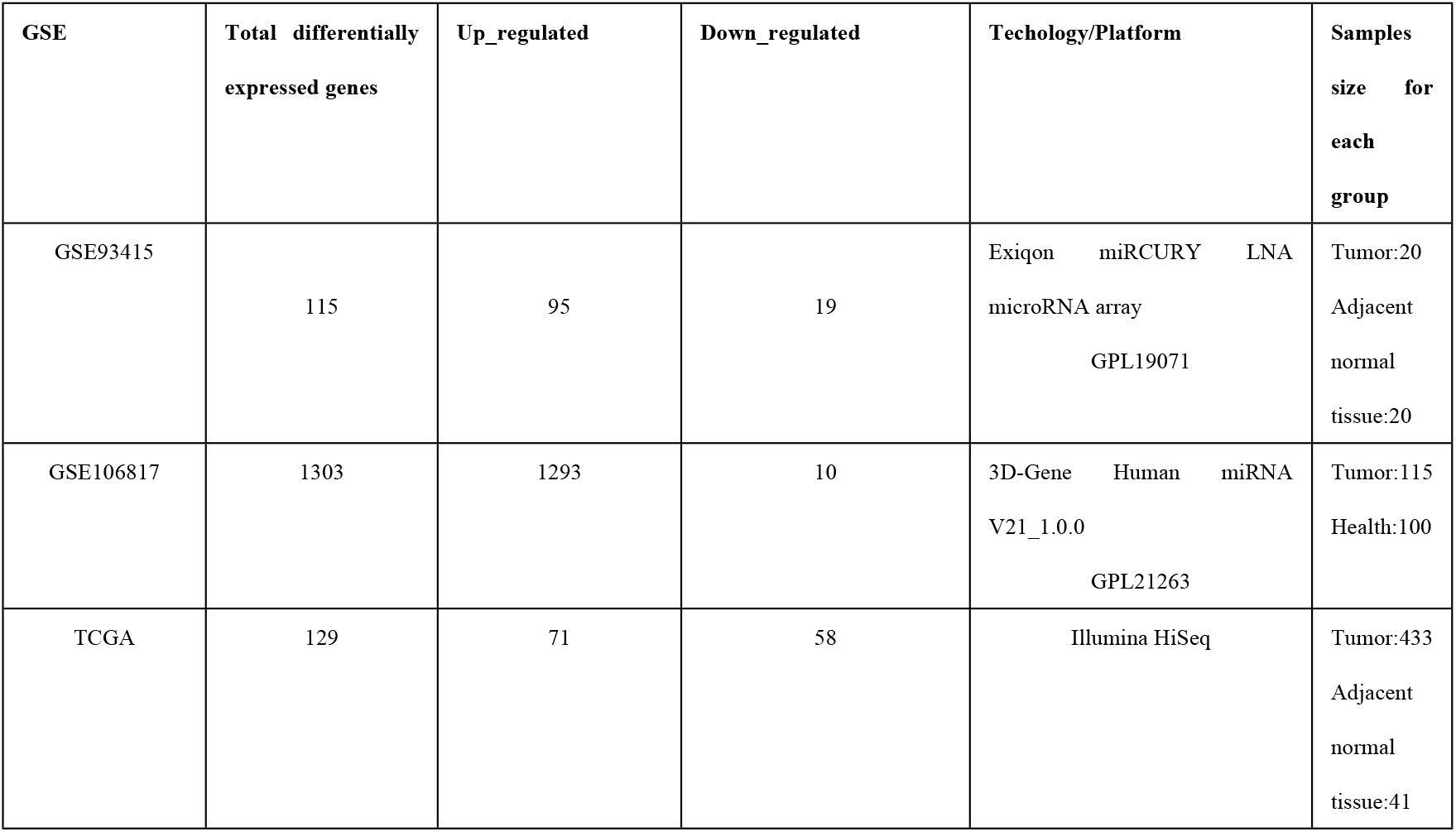
Details of GC studies and associated microarray datasets from GEO and TCGA database.

| GSE | Total differentially expressed genes | Up_regulated | Down_regulated | Techology/Platform | Samples size for each group |
| --- | --- | --- | --- | --- | --- |
| GSE93415 | 115 | 95 | 19 | Exiqon miRCURY LNA microRNA array<br>GPL19071 | Tumor:20<br>Adjacent normal tissue:20 |
| GSE106817 | 1303 | 1293 | 10 | 3D-Gene Human miRNA V21_1.0.0<br>GPL21263 | Tumor:115<br>Health:100 |
| TCGA | 129 | 71 | 58 | Illumina HiSeq | Tumor:433<br>Adjacent normal tissue:41 |

**Table 2.** Te clinical information and sample size for TCGA dataset.

|  |  |  |  |
| --- | --- | --- | --- |
|  | Alive = 267 | Dead = 176 | Total = 443 |
| Gender |  |  |  |
| Female | 100 (37.4%) | 59(33.5%) | 159(35.9%) |
| Male | 167 (62.6%) | 117(66.5%) | 284(64.1%) |
| Age |  |  |  |
| Mean(SD) | 64.9(11.10) | 66.7(10.17) | 65.6(10.8) |
| Median(Min,Max) | 67[30,90] | 66[41,87] | 67[30,90] |
| Stage |  |  |  |
| Stage I | 44(17.6%) | 14(8.5%) | 58 (13.9%) |
| Stage II | 92(36.6%) | 38(23%) | 131(31.2%) |
| Stage III | 97(38.7%) | 86(52.4%) | 183(43.9%) |
| Stage IV | 18(7.1%) | 26(15.8%) | 44(11%) |

**Figure 1.**
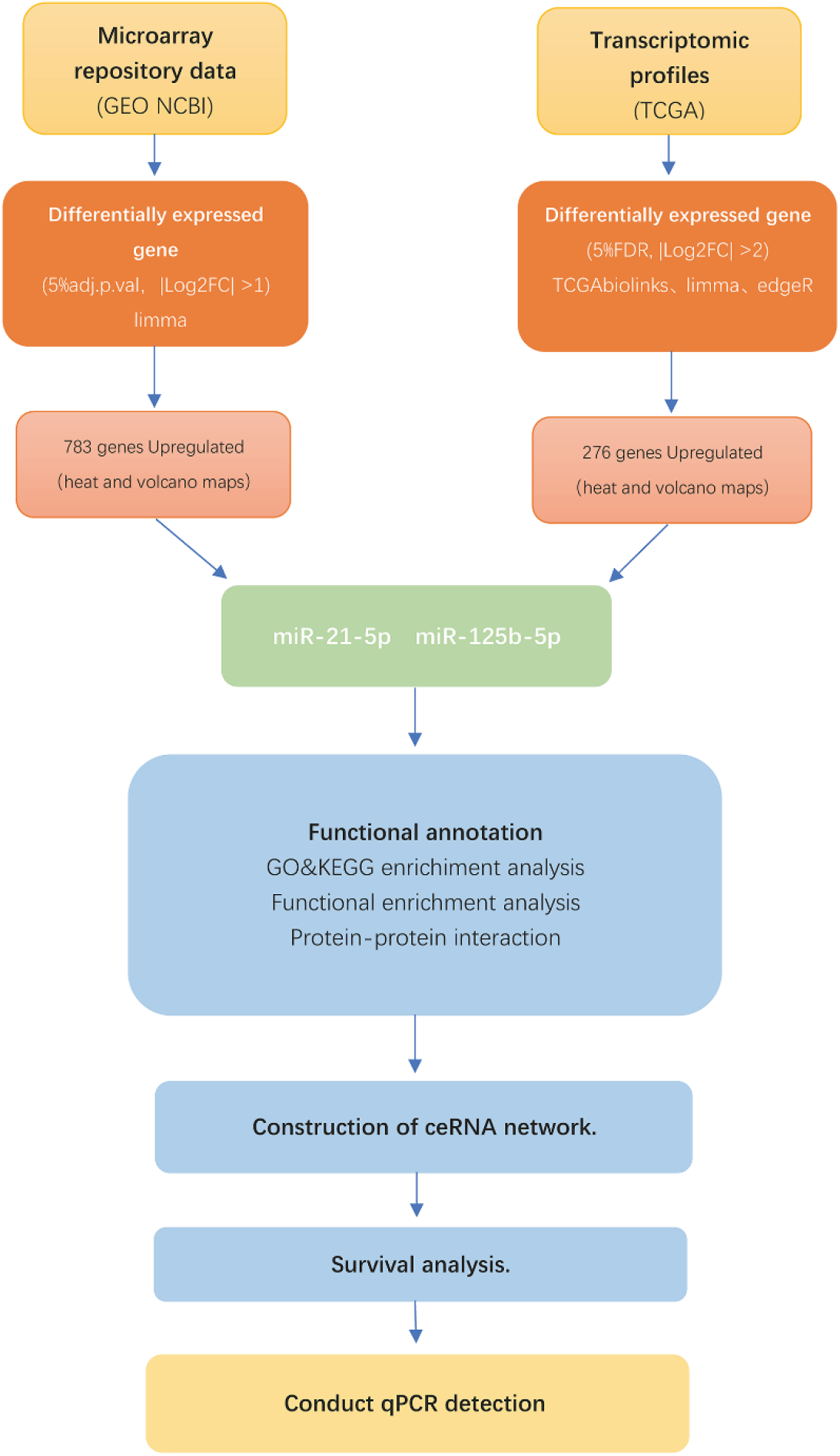
Flowchart for bioinformatics analysis of publicly available data from both GEO and TCGA databases.

### 2. Data processing and differential expressed miRNAs analysis in TCGA and GEO database

Using the Limma package in R software (version: x64 3.2.1), we performed background correction, standardization, and expression value calculation on the original datasets GSE93415 and GSE106817 from GEO. Differentially expressed miRNAs were screened based on fold-change (FC), adjusted p-values, and criteria of |Log (FC)|≥1 and adj. p-val<0.05. Similarly, miRNA expression quantification data and clinical information of GC patients from the TCGA database were downloaded using TCGAbiolinks, Limma, and edgeR to obtain a consensus of differentially expressed miRNAs. Parameters set for differential expression analysis were FDR<0.05 with |Log (FC)|≥1. Heatmaps and volcano plots were generated using the gplots package in R software. Subsequently, the intersected differentially expressed miRNAs from the two datasets were identified using the Venn package in R software.

### 3. Function Enrichment Analysis, GO and KEGG pathway analysis, and protein-protein interaction

Function enrichment analysis and GO and KEGG pathway analyses were conducted in R using the miPath V.3. Visualization of GO and KEGG was performed using GGPlot2. Protein-protein interaction analysis was performed using the STRING database (https://string-db.org/) and visualized using Cytoscape.

### 4. Construction of ceRNA network

We built a competing endogenous RNA (ceRNA) network by using GDCRNATools 16 to find if these two miRNAs competing endogenous regulating network. The major criteria of building ceRNA network in GDCRNATools are:(1) Expression of lncRNA and mRNA must be related. (2) The lncRNA and mRNA must share a great number of miRNA-21-5p, miRNA-125b-5p. Following the pipeline of GDCRNATools, we identified ceRNAs with the limma method (FDR<0.05 with |LogFC|≥1). The network was constructed by function of GDCRNATools. StarBase v2.0 was used to predict miRNA-mRNA interactions. Visualization of the ceRNA was performed by Cytoscape.

### 5. Survival analysis

To investigate the prognostic significance of the two identified miRNAs, survival analysis was conducted in R using Survtype. Clinical information was used to plot survival curves for patients with the relevant genes (*P*<0.05).

### 6. Conduct qPCR detection in SGC-7901, HGC-27 and GES-1

To validate our analysis results, qPCR was performed in SGC-7901, HGC-27, and GES-1 cell lines. These cell lines were obtained from the Type Culture Collection of the Chinese Academy of Sciences (Shanghai, China) and maintained in Dulbecco’s modified Eagle’s medium (DMEM) supplemented with 10%-20% fetal bovine serum (FBS) in a humidified cell culture incubator at 37°C and 5% CO2. Single-stranded cDNA was synthesized using the SuperScript First-Strand Synthesis System (Life Technologies, Carlsbad, CA, USA) following the manufacturer’s instructions. Primer sequences for miR-125b-5p and miR-21-5p were CCTCCCTGAGACCCTAACTTGTGA and CGCCGTAGCTTATCAGACTGATGTTGA, respectively. Real-time PCR reactions were performed using an ABI PRISM 7500 sequence detection system.

## Results

### 1. Identification of differentially expressed microRNAs and PCR

To distinguish the significant differences between normal and tumor samples in the GEO datasets, PCA was performed to reduce dimensionality and evaluate the independence of each group. The results demonstrated a significant difference between normal and tumor samples in the datasets (GSE106817, GSE93415) (Figure 2). The volcano plots illustrated significant differences between cancer and non-cancer samples in both datasets (Figure 3), with up-regulated and down-regulated miRNAs highlighted (Figure 4). Heatmaps of differentially expressed miRNAs in the GEO and TCGA datasets depicted their expression patterns in cancer and normal tissues (Figure 5A, B). The Venn diagram revealed 300 miRNAs present in both TCGA and GEO datasets (GSE106817, GSE93415) simultaneously (Figure 5C). Subsequently, 12 upregulated miRNAs in these datasets captured our attention. Finally, miRNA-21-5p and miRNA-125b-5p were selected for further research (Figure 6). The relative expression of miR-125-5p and miR-21-5p in SGC7901, HGC27, and GES-1 cell lines was determined by qPCR. Relative quantification of miRNA expression was calculated using the 2-ΔΔCt method, which represents relative fold changes in miRNA expression. Both miRNAs were found to be overexpressed in cancer cell lines, especially in SGC7901 (Figure 7).

**Figure 2.**
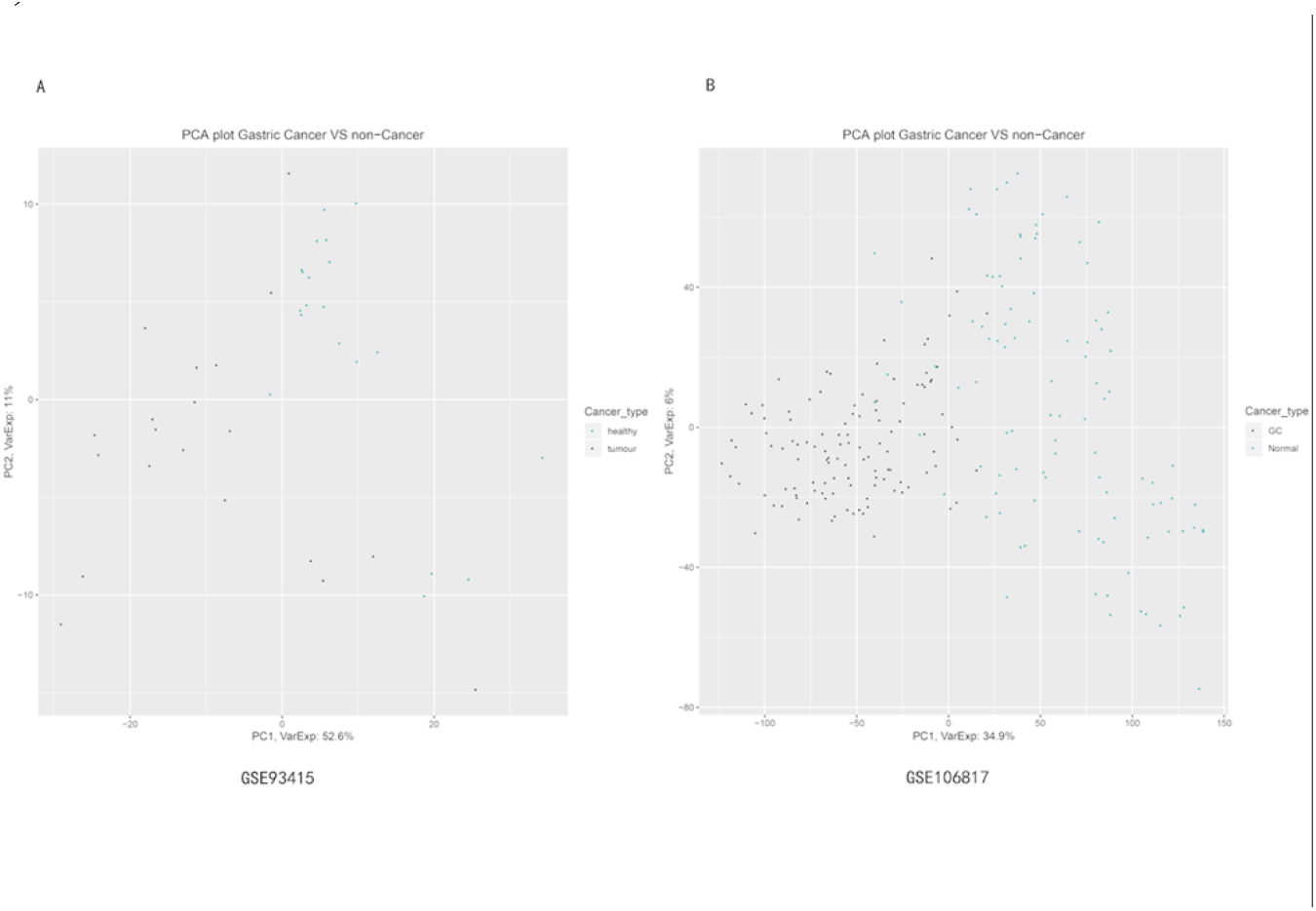
(A)The result of normal samples vs. tumor samples in GSE93415 displayed a significant difference. (B)The result of normal samples vs. tumor samples in GSE106817 displayed a significant difference.

**Figure 3.**
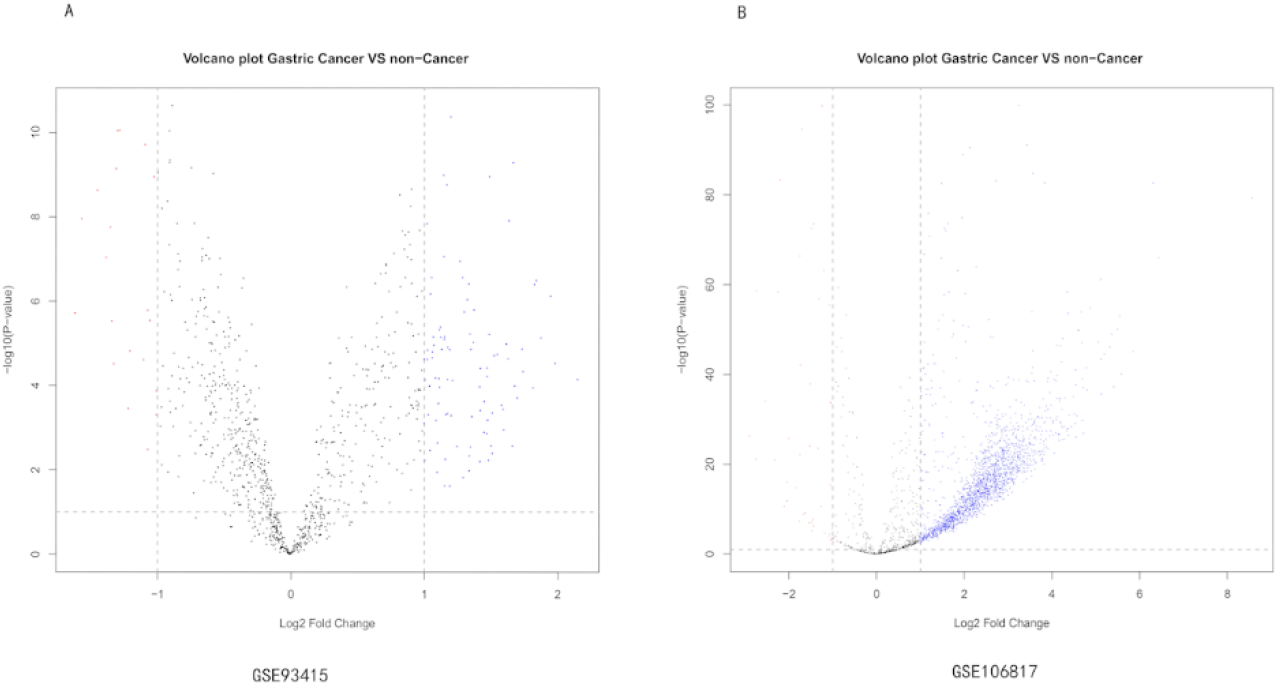
(A)The result of cancer vs. non-cancer in volcano plot from GSE93415. (B) The result of cancer vs. non-cancer in volcano plot from GSE106817.

**Figure 4.**
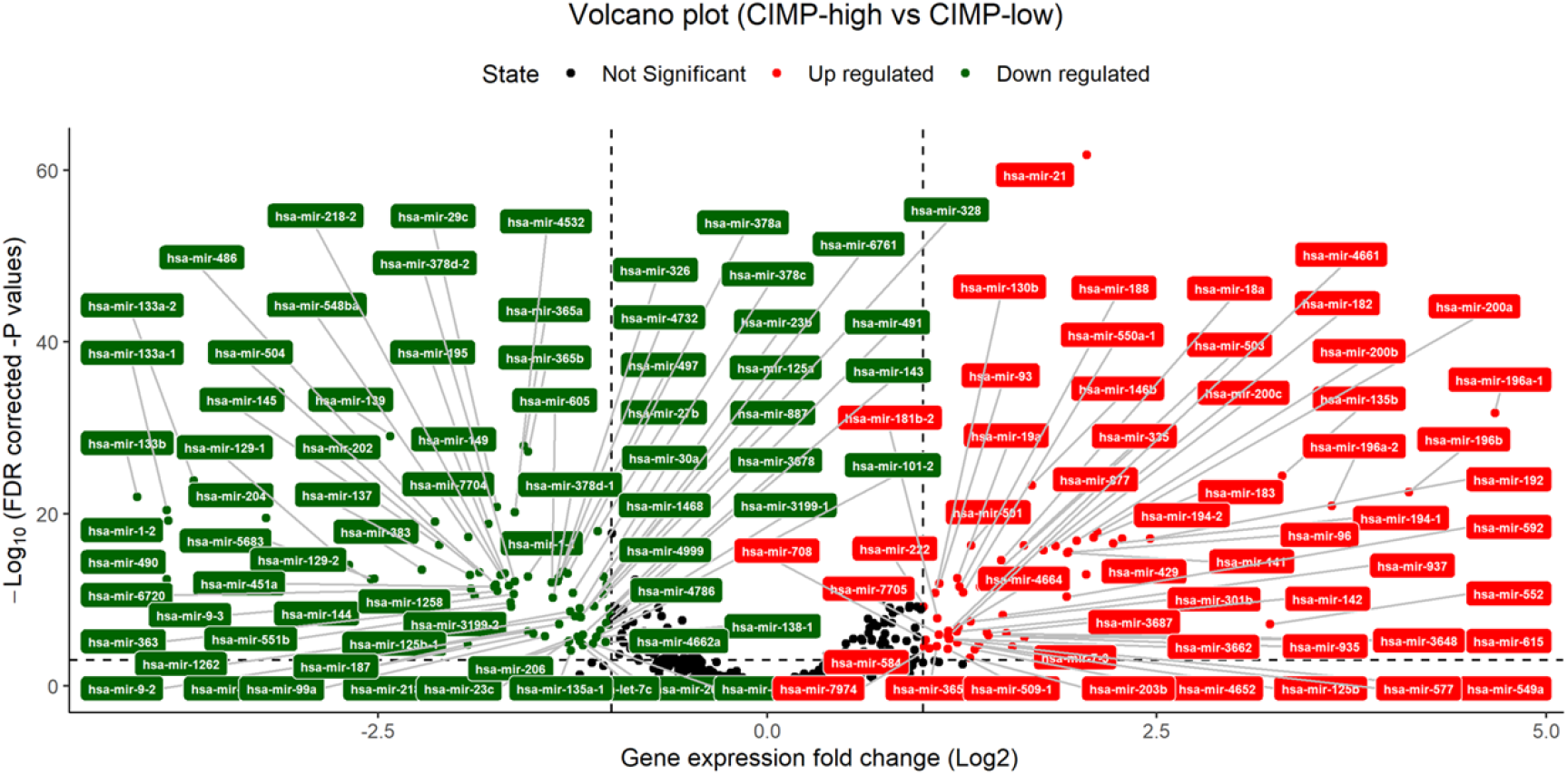
Volcano plot showed the number of differentially expressed genes in TCGA.

**Figure 5.**
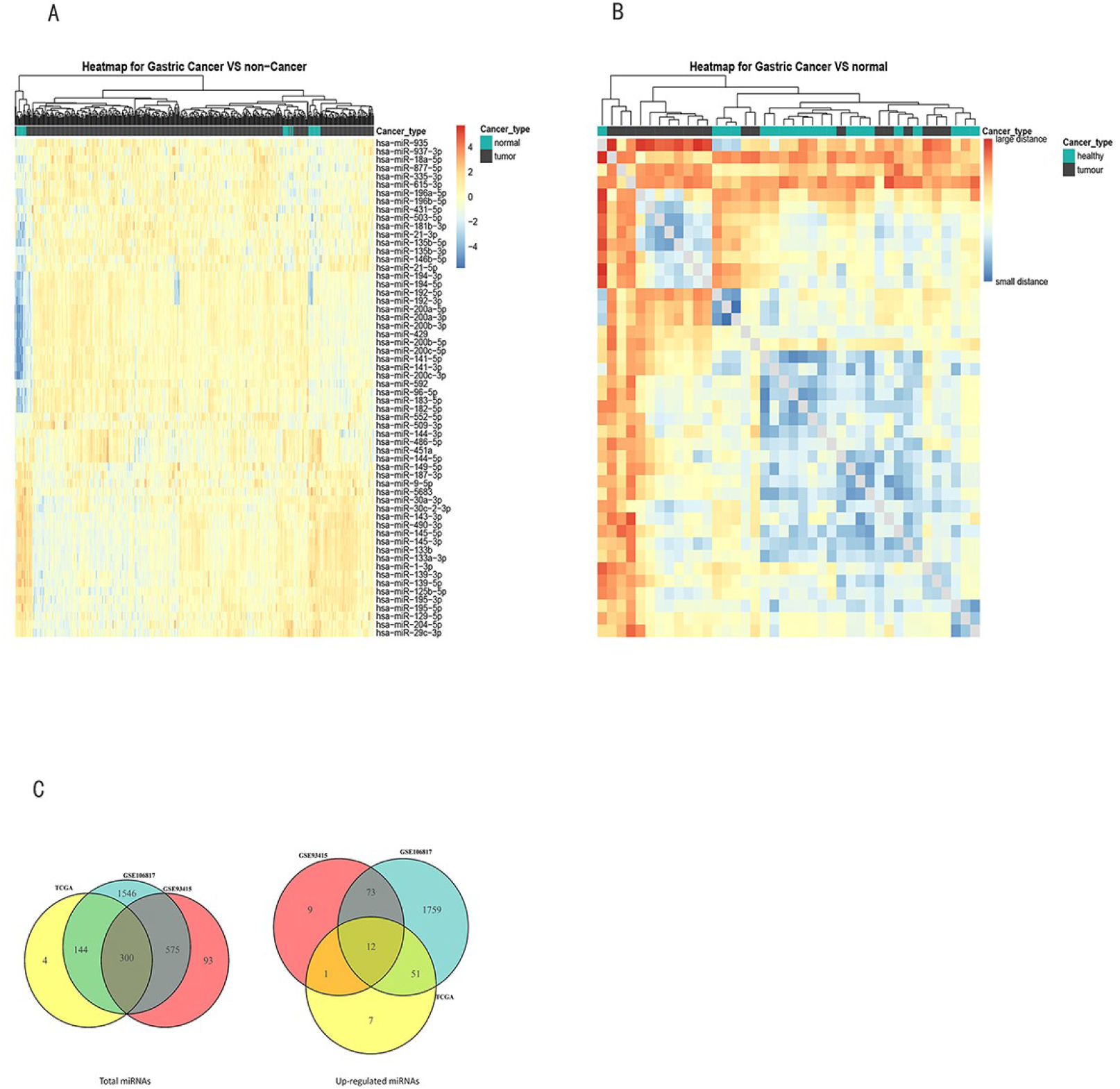
The expression changes of these genes in GEO and TCGA data. (A) Heatmap of differentially expressed genes in TCGA dataset coloring the samples-groups. (B) Heatmap of differentially expressed genes in GEO dataset coloring the groups. (C)Venn diagram demonstrates the intersections of genes between GEO data and TCGA data.

**Figure 6.**
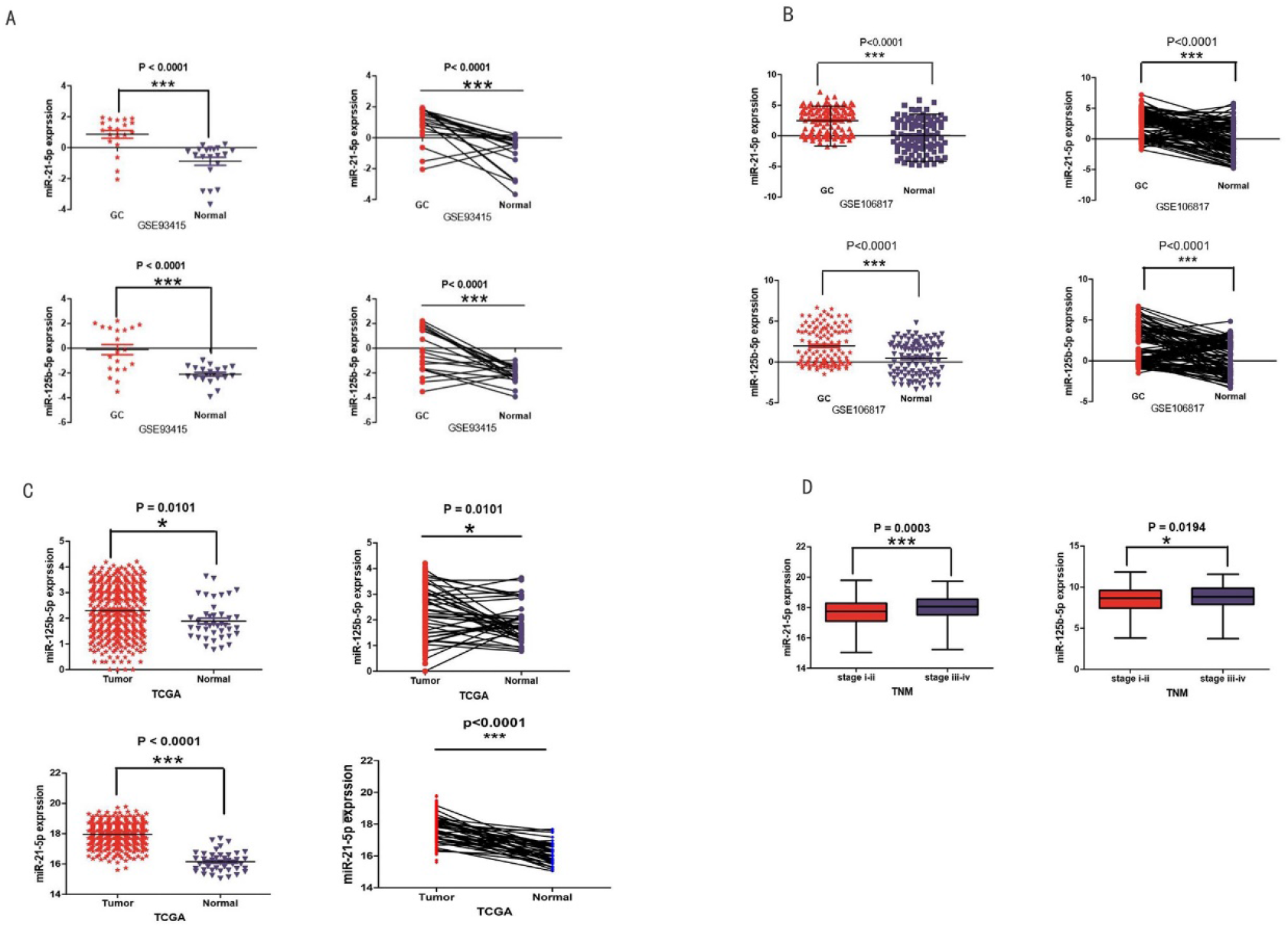
The expression of miRNAs in tumors and adjacent tissues in TCGA and GEO.(A)The expression of miR-21-5p and miR-125-5p in GSE93415. (B)The expression of miR-21-5p and miR-125-5p in GSE106817. (C)The expression of miR-21-5p and miR-125-5p in TCGA. (D) The expression of miR-21-5p and miR-125-5p in different TNM stage.

**Figure 7.**
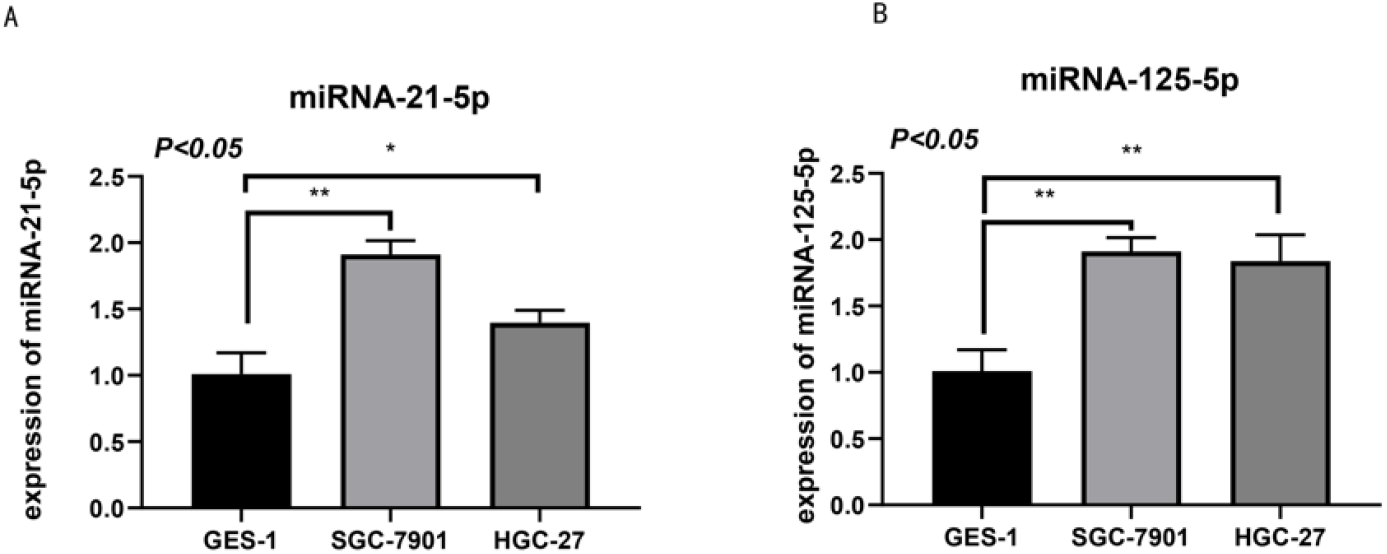
The expression of miR-125-5p and miR-21-5p in SGC7901, HGC27 and GES-1 cell lines in qPCR.

### 2. Survival analysis, GO terms and KEGG pathway analysis, functional enrichment analysis and protein-protein interaction

3. Based on TCGA and clinical information, we analyzed the survival of GC patients (Figure 8A). GO annotation results were categorized into three parts: biological process, molecular function, and cellular component (Figure 8B). The results revealed that the biological processes primarily associated with the two miRNAs were the mitotic cell cycle, viral processes, and stress-activated MAPK cascade. Additionally, cellular components exhibited a strong relationship with cell cellular components. According to molecular function, enzyme binding and protein binding transcription factor activity showed high correlations. Regarding KEGG pathway analysis, steroid biosynthesis was significantly enriched in the cell cycle. The STRING database (http://string-db.org) provided critical assessment and integration of protein interactions (Figure 8B). Relevant PPIs were obtained and visualized, revealing 48 nodes and 71 edges. Subsequently, we identified 8 stable and highest-scored genes (VPS4B, UBE2R2, MBOAT2, LVLAT1, BMF, SCARB2, TNFSF4, DIS3L2) in the network as hub genes (Figure 8C). Additionally, we analyzed the expression of 6 genes (VPS4B, UBE2R2, MBOAT2, LVLAT1, SCARB2, DIS3L2) in GC tumors and adjacent tissues (Figure 9A).

**Figure 8.**
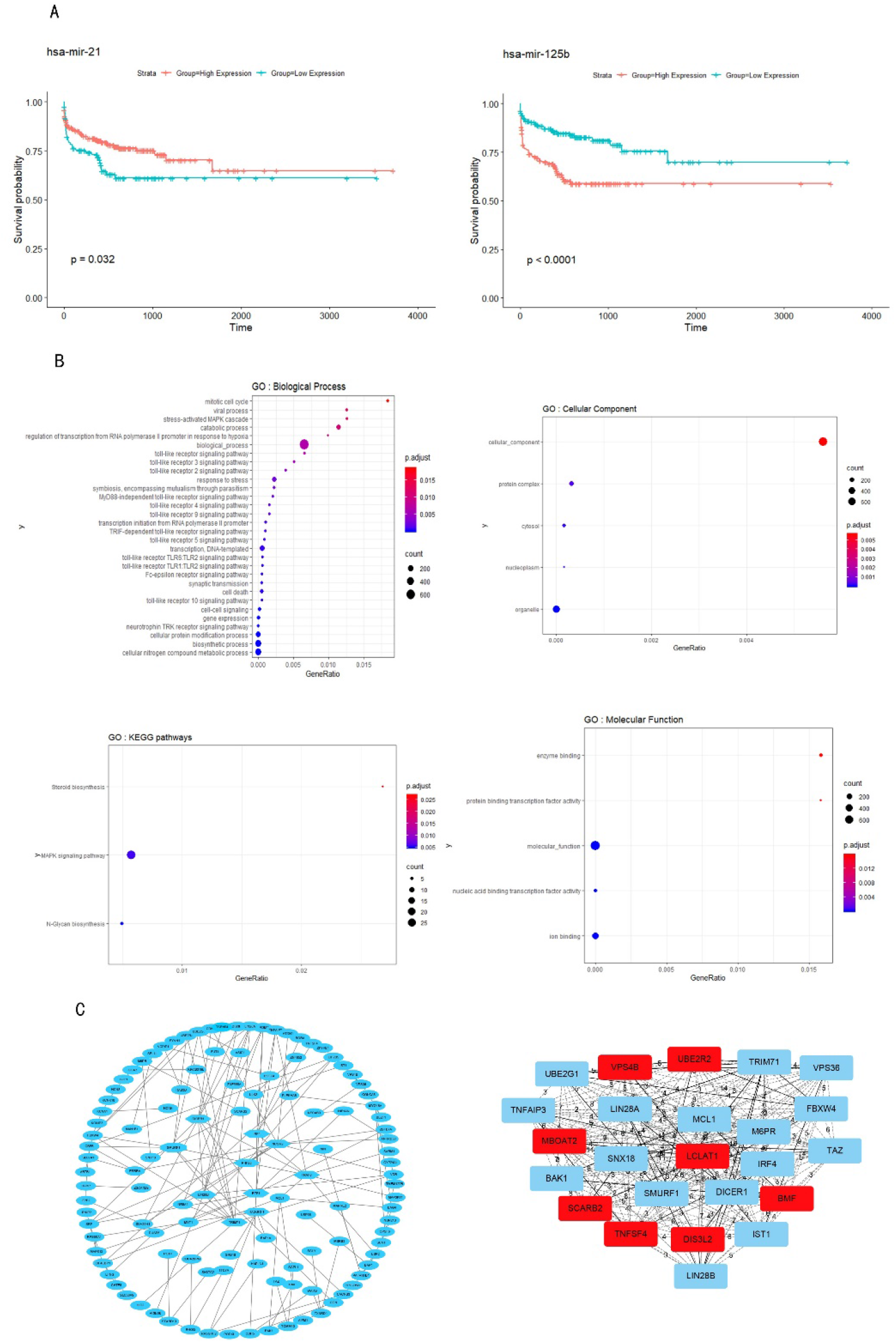
Survival analysis, GO terms and KEGG pathway analysis, functional enrichment analysis and protein-protein interaction in GC. (A) Survival analysis for differentially expressed genes in GC. (B) GO terms and KEGG pathway analysis, functional enrichment analysis in GC. (C) Protein-protein interaction network.

**Figure 9.**
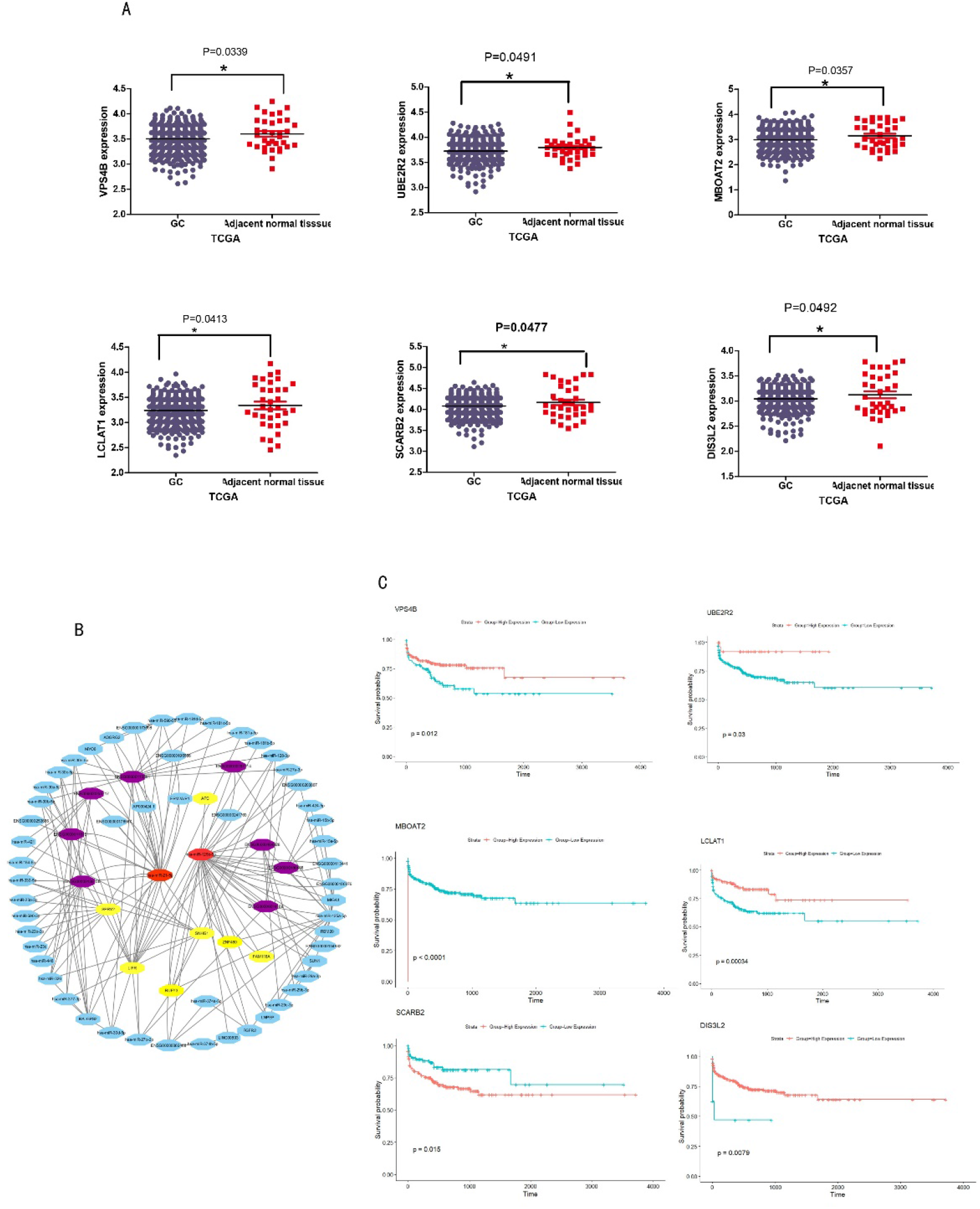
6 Hub genes (VPS4B, DIA3L2, LCLAT1, MBOAT2, SCARB2, UBE2R2) expression in GC, ceRNA network and survival analysis. (A) 6 hub genes expression in GC. (B) LncRNA– miRNA–mRNA interactions in GC. (C) 6 survival curves about the significance related (P<0.05) Hub genes.

### 3. ceRNA network and survival analysis

We identified 30 DElncRNAs (|LogFC|≥1, FDR<0.05) and 16 DEmRNAs (|LogFC|≥1, FDR<0.05) using GDCRNAtools, which exhibited significant differences in GC compared with controls.

Subsequently, miRNA-mRNA interactions were collected based on StarBase v2.0, and lncRNA-miRNA interactions were predicted based on miRcode. Visualization of the ceRNA network was performed using Cytoscape. Eight DElncRNAs (purple) and seven DEmRNAs (yellow) showed significant interactions with miRNA-21-5p and miRNA-125b-5p (Figure 9B).

Clinical information was obtained from TCGA. Based on TCGA data, we analyzed the survival curves of six hub genes (VPS4B, DIA3L2, LCLAT1, MBOAT2, SCARB2, UBE2R2) with significance (P<0.05). These six genes showed high expression in adjacent normal tissues, except for UBE2R2. The overexpression of VPS4B, DIA3L2, LCLAT1, MBOAT2, SCARB2 was associated with shorter survival times (Figure 9C).

## Discussion

Although advancements in surgical and medical treatments have provided hope for patients with gastric cancer (GC), the overall mortality rate has remained high over the past three decades[15]. The molecular mechanisms underlying GC are still unclear, and its morbidity and mortality rates continue to rank among the highest for cancer-related diseases[16-18]. Despite the significant improvements in early detection offered by gastroscopy, it is not a universally convenient method for early cancer detection, leading to a considerable number of gastric cancer patients being diagnosed at advanced stages[19]. Therefore, improving survival rates and prognosis remains a significant challenge. Exploring the molecular mechanisms of GC may help address this challenge. With the rapid development of microarray technology, the identification of general genetic alterations in malignant diseases, such as GC, may offer new insights[20, 21]. Additionally, recent technological advances in next-generation sequencing enable a more comprehensive and accurate examination of global gene expression profiles.

To identify potential biomarkers for GC prognosis and therapy, we utilized data from TCGA and GEO to access valuable information on GC. We discovered 12 up-regulated miRNAs that may be significantly correlated with GC. Among these miRNAs, miRNA-21-5p and miRNA-125b-5p caught our attention, prompting a detailed discussion of their functions. To further elucidate their underlying functions, we conducted functional enrichment analysis based on GO and pathway enrichment analysis based on KEGG. These upregulated miRNAs were primarily enriched in GO terms and pathways related to the mitotic cell cycle, viral processes, stress-activated MAPK cascade, and enzyme binding, which may contribute to tumorigenesis and tumor progression. Additionally, we identified a PPI network involving 8 important genes, namely VPS4B, UBE2R2,MBOAT2, LVLAT1,BMF,SCARB2,TNFSF4, and DIS3L2. Survival analysis based on these genes was also performed. Based on the above discussion, we analyzed that miRNA-21-5p and miRNA-125b-5p may improve survival time and serve as potential new targets for treatment.

miR-21-5p, encoded by the MIR21 gene located on chromosome 17q23.2 in humans[22], has gained prominence due to its significance in tumor progression and metastasis, particularly in cell proliferation and differentiation processes. Its expression is markedly increased in various solid tumors, including lung, breast, colon, gastric, and pancreatic cancer[23-28]. Furthermore, previous studies have identified miR-21-5p playing a crucial role in immune cells, promoting immune-related inflammatory diseases and autoimmune disease pathogenesis[29-32]. In our study, we observed upregulation of miRNA-21-5p in GC, suggesting that high expression of miRNA-21-5p may be associated with longer survival times. Additionally, downstream genes regulated by miRNA-21-5p may positively influence GC prognosis.

The function of miRNA-125b is very complicated, which was found to be abnormally expressed in several kinds of cancers., and function of it could as a carcinogen or tumor suppressor[33]. Previous studies have reported dysregulation of miRNA-125b-5p in cancers such as non-small-cell lung cancer[34] and cervical cancer[35]. Moreover, miRNA-125b-5p has been identified as tumor suppressor in various cancers such as in breast cancer[36]. Zhao et al. demonstrated that miRNA-125b-5p inhibits proliferation and migration by targeting SphK1 in bladder cancer. Thus, miRNA-125b-5p has been validated as a potential therapeutic target for cancer treatment and a biomarker. In our study, we observed upregulation of miRNA-125b-5p in GC, suggesting that high expression of miRNA-125-5p may be associated with shorter survival times. Additionally, downstream genes regulated by miRNA-125b-5p may also influence GC prognosis positively. However, the precise functions and mechanisms of these two miRNAs in GC remain unclear.

## Conclusion

In summary, our study analyzed the array-based and sequence-based data of GC supported by GEO and TCGA databases. miRNA-125-5p and miRNA-21-5p were discussed by bioinformation in the above. Both of them have over expression in GC cancer than the adjacent tissues which may contribute to the finding of molecular mechanisms underlying the initiation and development of GC.

## Abbreviations

miRNA: MicroRNA
GEO: Gene Expression Omnibus
TCGA: Cancer Genome Atlas
GC: Gastric cancer
ceRNA: Competing endogenous RNA
DMEM: Dulbecco’s modified Eagle’s medium
FBS: Fetal bovine serum

